# A SuperSelective primer-based real-time PCR platform for hypersensitive detection of azole heteroresistance in *Cryptococcus neoformans*

**DOI:** 10.64898/2026.04.23.720376

**Authors:** Siddhi Pawar, Honglin H. Xue, Sophia Wang, Salvatore Marras, Chaoyang Xue

**Author notes:** Corresponding authors: Salvatore Marras, Chaoyang Xue.

## Abstract

*Cryptococcus neoformans* is the leading cause of fungal meningitis with limited treatment options, making early and accurate diagnosis critical for improved patient outcome. Current diagnostic methods for cryptococcosis rely largely on capsule antigen detection and fungal culture, which are time-consuming and unable to identify mutation-driven antifungal heteroresistance. In this study, we have developed a SuperSelective primer-based PCR (SSP-PCR) platform for the rapid and specific detection of azole resistance-associated single-nucleotide polymorphisms (SNPs). We demonstrate that SSP-PCR reliably detected a single copy mutant allele in the presence of excess wild-type (WT) DNA, with sensitivity reaching a 1:10^4^ mutant-to-WT ratio. Incorporating molecular beacon (MB) probes with our SSP-PCR platform further enhanced amplification specificity, enabling selective detection of the *ERG11* Y145F (A434T) mutation that is known to cause high azole resistance. Using genomic DNA from in vitro cultures and mouse lung tissues infected with either WT strain H99 strain or a fluconazole-hyper-resistant *mrl1* clinical isolate that carries the *ERG11*(A434T) mutation, or both strains, we successfully detected the A434T mutant allele in both settings. Moreover, our SSP-PCR simultaneously identified *ERG11*(A434T) and the multi-azole resistance-associated *ERG11*(G1885A) mutant alleles in a single-tube duplex reaction. Collectively, the SSP-PCR platform provides a robust and ultrasensitive molecular approach for the detection of azole resistance and heteroresistance in *C. neoformans*, with strong potential for high-throughput clinical screening applications.

**Importance:** Invasive fungal infections are a growing public health threat, causing over 1.5 million deaths annually, with cryptococcal meningitis accounting for over 15% HIV/AIDS related mortality. The problem is aggravated by limited treatment options and emerging of drug resistance. Long-term use of fungistatic azoles like fluconazole promotes the emergence of azole heteroresistance, contributing to clinical treatment failure. Current diagnostic assays often fail to detect resistance-associated mutations within heterogeneous fungal populations, limiting their clinical utility. In this study, we developed a SuperSelective primer-based PCR (SSP-PCR) platform for the rapid and ultrasensitive detection of azole resistance associated single-nucleotide polymorphisms (SNPs) in *Cryptococcus neoformans*. By integrating molecular beacon probes, this assay achieves high specificity and enables simultaneously detection of multiple SNP mutations in a single reaction. Our SSP-PCR platform offers a powerful molecular approach for identifying azole resistance and heteroresistance, with strong potential to improve diagnostic precision and guide antifungal therapy in clinical settings.

## Introduction

Invasive fungal infections represent an increasing public health threat worldwide, causing over 1.5 million deaths annually (1)(2)(3). *Cryptococcus neoformans* (*Cn*) is an opportunistic fungal pathogen that causes infection through inhalation of spores and yeasts. Following pulmonary infection, *Cn* can disseminate to the central nervous system to cause life-threatening fungal meningitis, accounting for ∼15% HIV/AIDS related deaths globally (4). Despite these alarming mortality rates, therapeutic options remain limited, and the problem is aggravated by their collateral effects and drug resistance. Currently, polyenes and azoles are the backbone of anticryptococcal therapy. The polyene Amphotericin B (AmB) is a rapidly acting fungicidal against *Cn*, yet highly toxic (5). Patients with cryptococcal meningitis are typically treated with AmB and 5-flucytosine first, then switched to triazoles such as fluconazole (FLC) for long-term maintenance therapy. FLC is the preferred choice for maintenance therapy due to its low toxicity and oral bioactivity. However, long-term use of fungistatic azoles like FLC has allowed the fungus to persist in immunocompromised hosts. The repeated exposure to FLC enables the fungal isolates to mutate and develop heteroresistance, resulting in suboptimal clinical outcomes (6, 7). Heteroresistance has been demonstrated as a mechanism of clinical treatment failure, without alterations in azole susceptibility, in cryptococcal meningitis patients on long-term FLC maintenance therapy (8)(9)(10)(11)(12). Compounding this challenge, the lack of simple diagnostic methods to identify potential azole resistance and heteroresistance in clinical samples contributes to poor patient outcomes.

The current diagnosis of cryptococcosis relies on a combination of histopathology, direct microscopy, fungal culture, and detection of cryptococcal antigen (CrAg) in bodily fluids. Among these, antigen-based assays are widely used for their high sensitivity and high specificity (13). These assays detect capsular polysaccharide antigen in serum and cerebrospinal fluid. In clinical settings, commonly used CrAg detection methods include enzyme linked immunosorbent assay (ELISA), latex agglutination, and lateral flow assay (LFA) (13). Of these, LFA is the most employed diagnostic approach in clinical settings because it is rapid, simple, and cost-effective. However, none of these methods can detect mutation-induced resistance or heteroresistance.

Azole resistance mostly occurs due to overexpression and/or mutations in the drug target, lanosterol 14-alpha-demethylase (*ERG11*) gene. A single nucleotide mutation in *ERG11* can confer resistance (14)(15). Extensive analysis of clinical and environmental isolates of *Cn* and *C. gattii* by Dr. Kwon-Chung’s group revealed that the Y145F (A434T) single mutation in *ERG11* causes high-level FLC resistance (9), while the *ERG11*(G1885A) point mutation results in multi-azole resistance (16). Detecting heteroresistance within a mixed fungal population remains extremely challenging and is not achievable with currently available diagnostic techniques, even with molecular methods such as PCR and whole-genome sequencing.

In this study, we have developed a SuperSelective primer-based real-time PCR (SSP-PCR) platform for rapid and specific detection of azole resistance associated with single-nucleotide polymorphisms (SNPs) in the *Cn ERG11* gene, including *ERG11*(A434T) and *ERG11*(G1885A). The unique design of SuperSelective primers enables highly specific recognition and quantitative detection of mutant targets, even in the presence of closely related wild-type sequences (17-19). Upon hybridization to the target sequence, these primers initiate synthesis of mutant DNA fragments, generating amplicons that undergo exponential amplification in subsequent thermal cycles (17-19). To our knowledge, this is the first study to employ SuperSelective primer design for the detection of azole-associated antifungal drug resistance mutation in *Cn*. Additionally, the incorporation of molecular beacons (MBs) enabled the development of a duplex real-time PCR assay that simultaneously detects two mutant target sequences in a single reaction. MBs are stem-loop-shaped oligonucleotide probes that fluoresce upon hybridization to DNA or RNA target sequences (20)(21) and have been widely used to identify different human pathogens, including fungi (22). Collectively, our work established the first SSP-PCR platform for the rapid detection of azole-resistance associated SNP mutations in *Cn*, validated in vitro using both synthetic g-blocks and genomic DNA and in vivo infected tissues.

## Results

### Designing SuperSelective primers for real-time PCR detection of *ERG11*-associated mutations

SuperSelective primers consist of three parts: (a) anchor sequence; (b) bridge sequence, and (c) foot sequence (Fig. 1A). The 5’-anchor sequence, like a conventional PCR primer, binds only to the DNA region of interest. The 3’-foot sequence is separated from the anchor by a bridge sequence that forms a single-stranded bubble in the resulting hybrid. The 3’-foot sequence is a short sequence, possesses an “interrogating nucleotide” allowing it to form complementary hybrids with mutation-containing target sequence, resulting in generation of amplicons (Fig. 1A). Following the design guidelines by Vargas et al (17), we designed SuperSelective primers for detection of *Cn ERG11*(A434T) and *Cn ERG11*(G1885A) mutations that are known to cause high-level FLC resistance (Table 1). The SuperSelective primer designation 20:14/15:6:1:0 of *ERG11*(G1885A) indicates a 20-nucleotide anchor, 14-nucleotide bridge opposite a 15-nucleotide intervening template sequence, and a 7-nucleotide foot composed of 6-nucleotides + 1 interrogating nucleotide positioned at the 3’ end. Similarly, the SuperSelective primer designation of *ERG11*(A434T) 24:13/14:8:1:0 indicates a 24-nucleotide anchor, 13-nucleotide bridge opposite a 14-nucleotide intervening template sequence, and a 9-nucleotide foot composed of 8-nucleotides +1 interrogating nucleotide positioned at 3’ end.

**Table 1.**
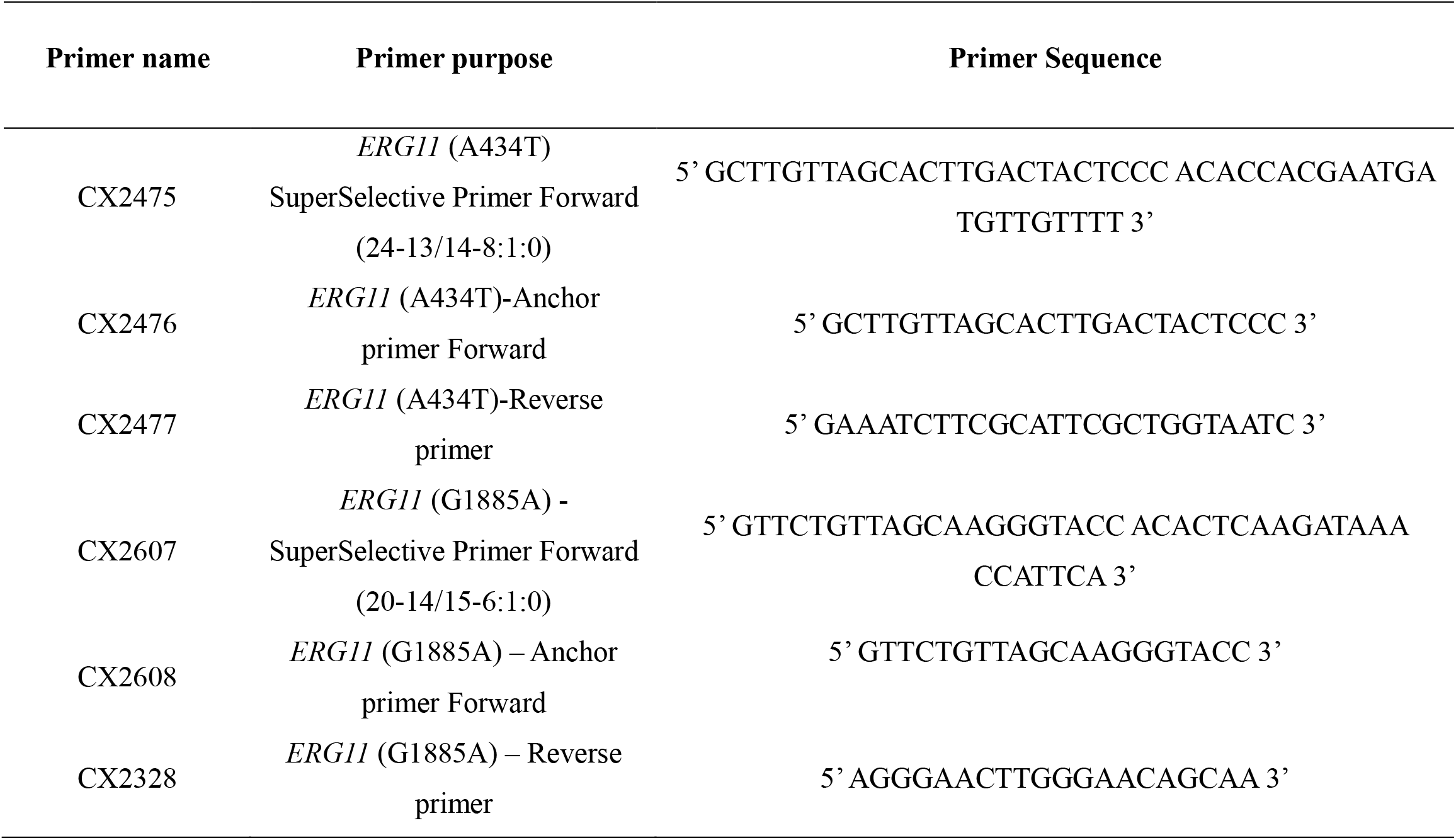
Primer sequences used in this study.

**Figure 1.**
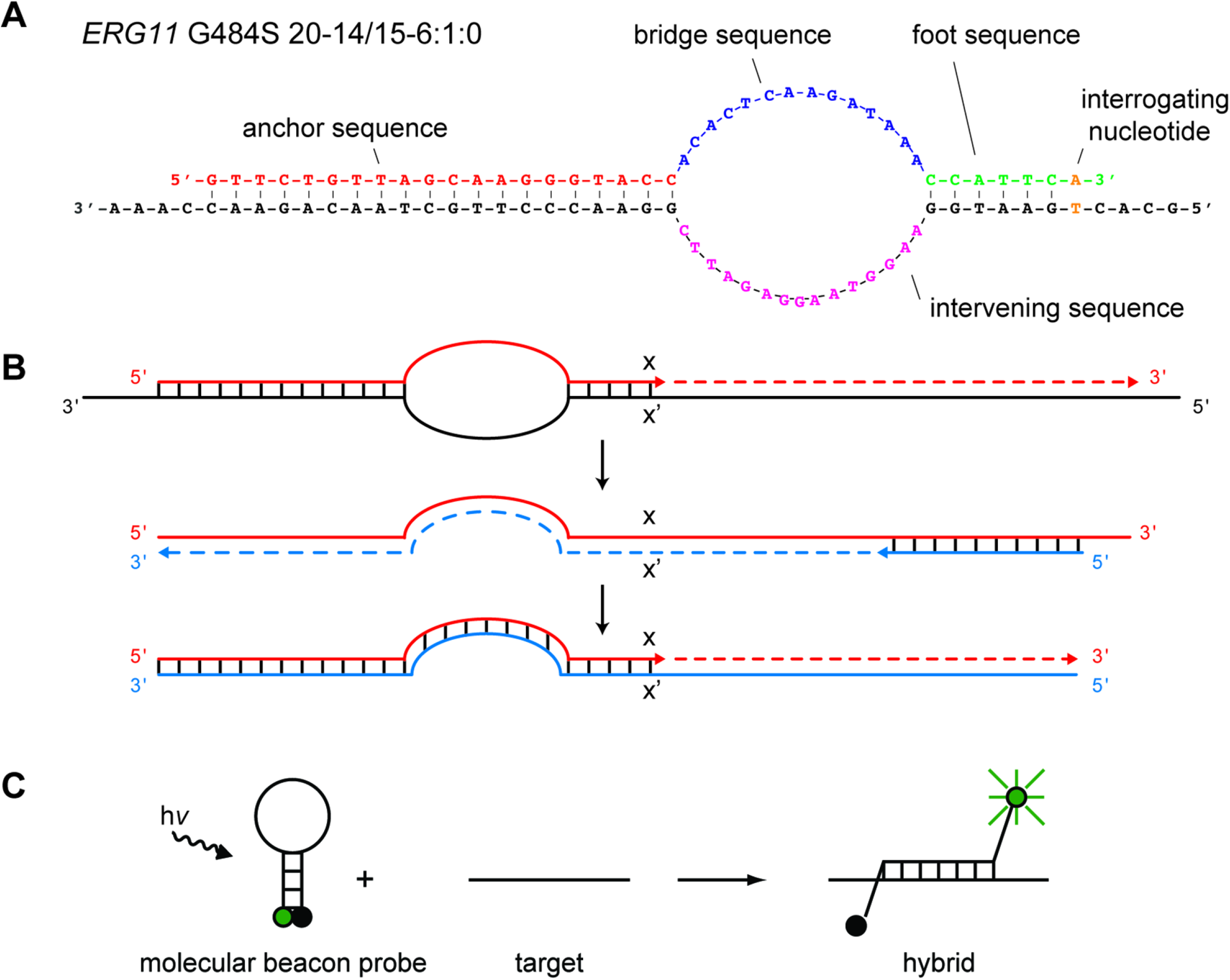
Schematic illustration of SuperSelective primers and molecular beacon design. (A) Structure of a SuperSelective primer for detecting *ERG11*(G1885A) mutant sequences in the presence of *ERG11* wild type sequences. (B) Principle of operation of SuperSelective primers I) hybridization of the SuperSelective primer with the mutant DNA template fragment, which depends on matching the anchor and the foot sequences (x and x’ denotes the SNP), leading to synthesis of a (+) amplicon (red line) from a (-) template. II) The (+) amplicon serves as a template for a conventional reverse primer, generating a (-) amplicon (blue-dotted line), with a unique bridge sequence that is complementary to the SuperSelective primer. III) In subsequent thermal cycles, the entire SuperSelective primer sequence is complementary to the (-) amplicon strand, leading to exponential amplification. (C) The principal operation of molecular beacons.

To monitor the PCR amplification signal in real time, we employed the MB probes that bind to the PCR amplicon. In this study, we designed MB probes that binds to *ERG11*(A434T) and *ERG11*(G1885A) regions of *Cn* DNA, respectively (Table 2).

**Table 2.**
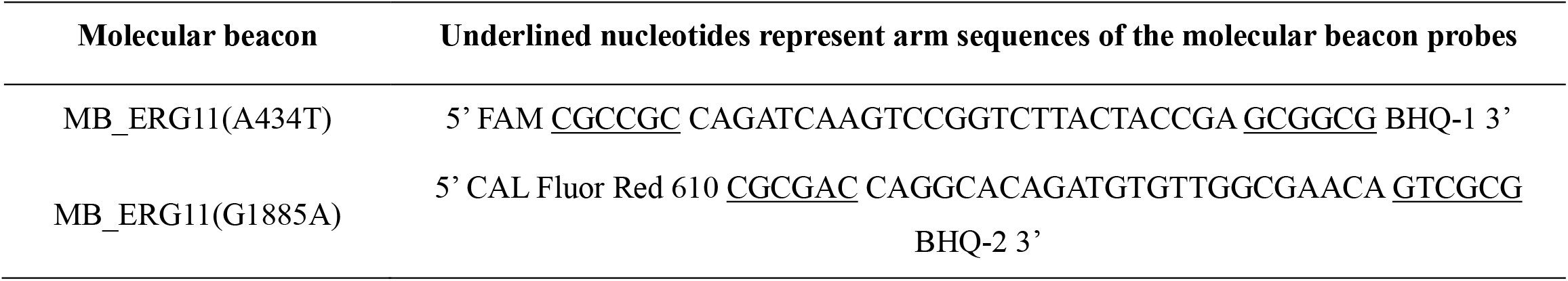
Molecular beacon design for *ERG11* Y145F(A434T) and G484S(G1885A) mutation detection.

### Determining the sensitivity and specificity of the SSP-PCR

We designed a real-time PCR to detect the SNP mutation in *ERG11*(A434T) (T<u>A</u>C->T<u>T</u>C) using a SuperSelective primer. To determine SSP-PCR specificity, we first used synthetic 400 bp g-Block templates (an ERG11 wild-type template and an *ERG11*(A434T) (T<u>A</u>C->T<u>T</u>C) template), allowing us to accurately calculate the template DNA copy number in each PCR assay (Table S1). Using the conventional primer set (CX2476 and CX2477) and SYBR Green to monitor the real-time PCR assay, we observed identical PCR threshold cycles (Ct) from both wild type *ERG11* and *ERG11* (A434T) mutant templates, confirming the same amount of DNA templates were used (Fig. 2A). Then, we performed real-time PCR assays with the SuperSelective primer CX2475 and the reverse primer CX2477 using the same amount of DNA as template. We found that the SuperSelective primer set amplified the *ERG11*(A434T) mutant DNA template with a significantly earlier Ct value of 15 for 10^6^ copies compared to the same amount wild type (*ERG11*WT) template that was amplified 20 cycles later with a Ct value of 35 (Fig. 2B). Furthermore, the SSP-PCR robustly amplified the *ERG11*(A434T) mutant template, yielding Ct values of 25 and 35 with 10^4^ and 10^2^ copies of DNA template, respectively. In contrast, no amplification was observed with WT templates at the same copy numbers (Fig. 2B).

**Figure 2.**
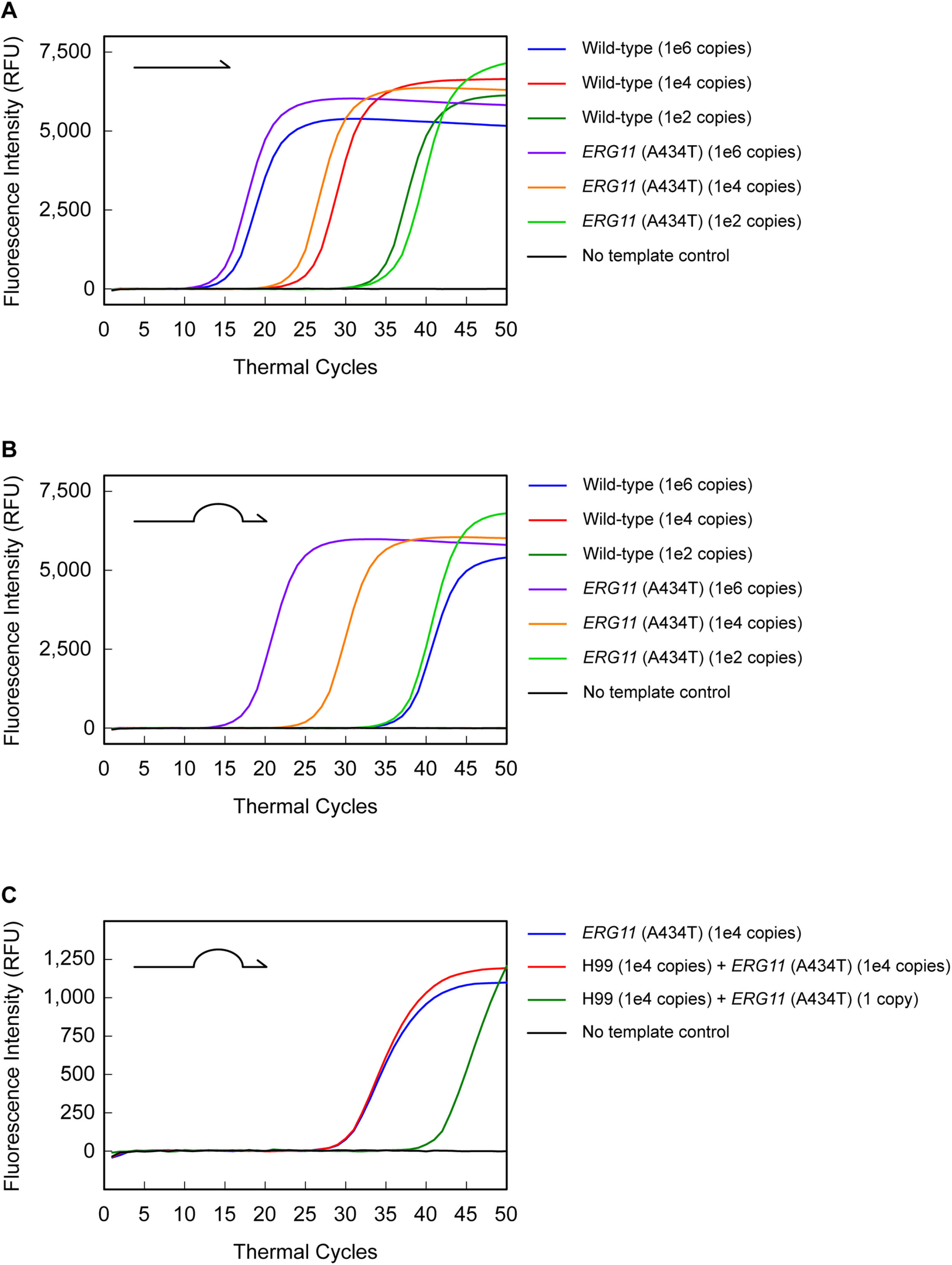
SSP-PCR assay detects *C. neoformans ERG11(*A434T) mutation with high specificity and sensitivity in synthetic g-block DNA. (A) The plot shows real-time PCR amplification curves generated with conventional primers using templates containing either the *ERG11*(A434T) mutation or the wild-type (WT) *ERG11* sequence. These two templates showed comparable amplification at equivalent input copy numbers of 10^6^, 10^4^, and 10^2^, confirming equal DNA copies across reactions. (B) Real-time PCR amplification curves obtained with SuperSelective primers using templates containing either the *ERG11* (A434T) mutation or the WT sequence. 10^6^, 10^4^, and 10^2^ copies of the mutant template were detected, whereas only the WT template (10^6^ copies) showed significant delayed amplification (∼Ct 35), demonstrating the specificity of the SuperSelective primers for *ERG11*(A434T). (C) The plot shows real-time amplification curves obtained in the SSP-PCR assay using *ERG11*(A434T) mutant templates in the presence of 10^4^ copies of the WT template. Robust detection of mutant templates in these conditions demonstrates the sensitivity of the assay for identifying low-abundance mutants in an excess WT background. No amplification observed in the no-template control (NTC).

To evaluate the sensitivity of SSP-PCR assays, we mixed wild-type (*ERG11*WT) and *ERG11*(A434T) mutant DNA templates at *ERG11*WT 10^4^: *ERG11(*A434T 10^4^) and *ERG11*WT 10^4^:1 *ERG11*(A434T) ratio and subjected them to real-time PCR using SuperSelective primers integrated with MB probe MB_ERG11(A434T) for their detection (Table 2). The SSP-PCR assay generated a fluorescence signal for the *ERG11*(A434T) mutant template in both DNA mixtures. SuperSelective primers specifically identified a single copy of the *ERG11*(A434T) mutation DNA even in the presence of 10^4^ copies of *the ERG11*WT template (Fig. 2C), demonstrating the sensitivity of the SSP-PCR. No WT template amplification was detected, indicating a high PCR specificity. Together, these results confirmed that our SSP-PCR platform enables a robust detection of SNP variants in the presence of abundant WT templates.

### Detection of *ERG11*(A434T) SNP signal in genomic DNA of clinical isolates from in vitro cultures and in vivo infected mouse lungs

To determine the potential application of the SSP-PCR platform as a future diagnostic method, we tested the PCR assay using genomic DNA from clinical isolates, including the reference strain H99 and an *mrl1* strain carrying the *ERG11*(A434T) mutation. We isolated genomic DNA from fungal strains grown on YPD medium and successfully detected the *ERG11*(A434T) mutation in a PCR assay with 100 ng template DNA in each reaction (Fig. 3A). Using conventional primers CX2476/CX2477, the PCR with both templates had a Ct at ∼20 cycles, indicating an equal amount of the template DNA in each reaction (Fig. 3A). In our SSP-PCR assay with SuperSelective primer CX2475 and the reverse primer CX2477, we detected the MB signal at 20 cycles with the *mrl1* DNA, while a later Ct of 35 cycles was observed with the H99 template (Fig. 3A), indicating a high *ERG11*(A434T) SNP detection specificity.

**Figure 3.**
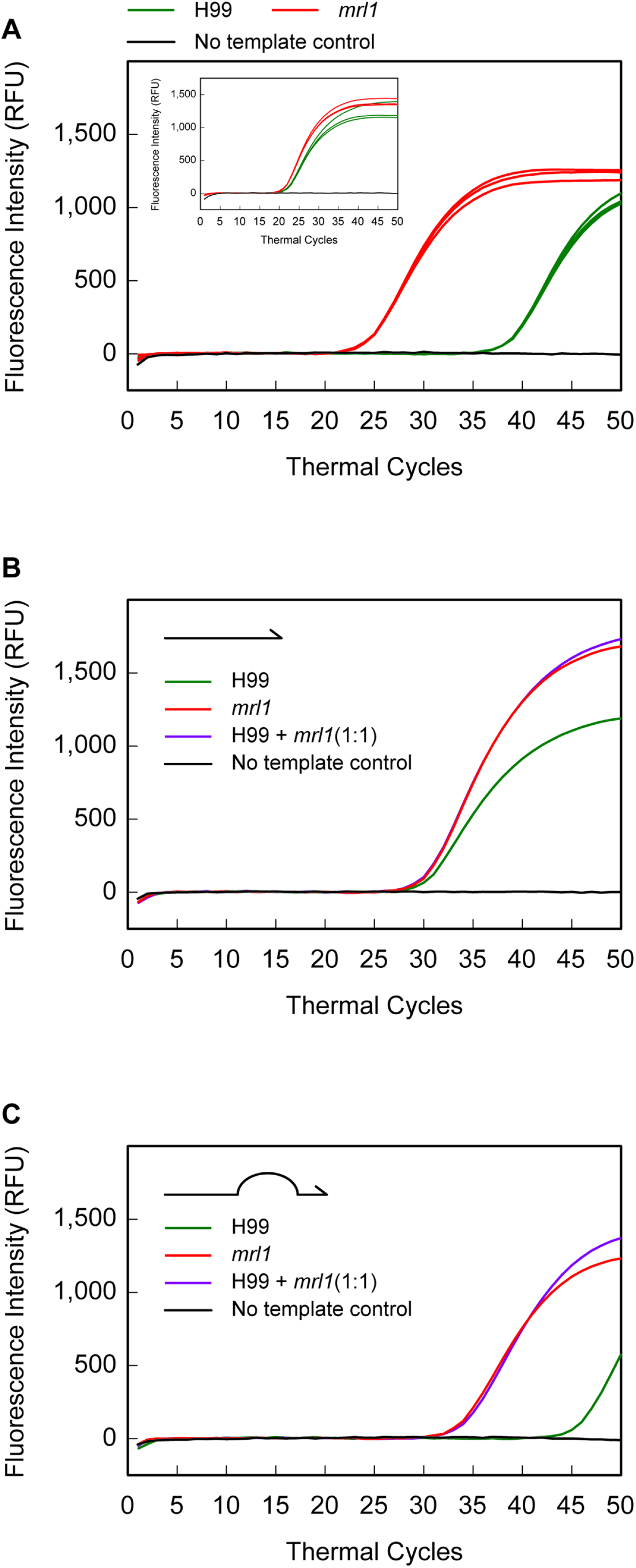
Detection of *C. neoformans* and azole-associated *ERG11*(A434T) mutation in genomic DNA. (A) The *ERG11*(A434T)-specific molecular beacon selectively detected the azole-resistance-associated mutation in genomic DNA isolated from *the C. neoformans ERG11* (A434T) mutant strain (*mrl1)* at an earlier Ct (∼20 cycles) than in WT H99 genomic DNA. The insert demonstrates comparable DNA input across samples (Ct ∼20 for all samples), as confirmed by equal amplification with the conventional primers. In vivo detection of *C. neoformans* from mouse lungs infected with 10^5^ H99, 10^5^ *mrl1* mutation, or mixed H99 and *mrl1* cells at a 1:1 ratio. (B) Real-time amplification curves generated utilizing the conventional primer set show that all reactions contain approximately the same total number of DNA templates, suggesting similar fungal burden in each sample. (C) The plot shows real-time amplification curves for *mrl1* mutation at approximately Ct∼20, confirming sensitive detection of *mrl1* in vivo infected tissue samples.

Furthermore, to determine the sensitivity of this detection method *in vivo* in infected tissues, we intranasally infected the C57BL/6 mice with 10^4^ cells of either H99 only, a *mrl* mutant strain only, or a mixed inoculum containing both strains at 1:1 ratio (10^4^:10^4^). Infected lung tissues were isolated at day 7 post-infection, and their total genomic DNA was extracted using the CTAB method. Using 45 ng of these total genomic DNAs as templates, our PCR with conventional primer CX2476 /CX2477 resulted in identical Cts (∼30 cycles), indicating an equal number of fungal DNA copies for each reaction (Fig. 3B). We then performed our SSP-PCR assay using primers CX2475 and the reverse primer CX2477. Our MB probe successfully detected the *ERG11* (A434T) mutation in at much earlier PCR cycles at 30-35 cycles than the wild type, which was not amplified until over 45 cycles, indicating a high amplification specificity (Fig. 3C). Overall, our results demonstrate that our SSP-PCR assay in combination with MB probe, can detect the SNP mutations in genomic DNAs isolated from *C. neoformans* strains or their infected lungs.

### Duplex detection of *ERG11* (A434T) and *ERG11* (G1885A) mutations

Clinical specimens may harbor mixed *Cryptococcus* populations that contain multiple azole-resistance alleles. To assess whether our SSP-PCR platform can detect two mutations simultaneously, we developed a duplex assay as a proof-of-concept. In addition to the *ERG11* (A434T) mutation, we designed a SuperSelective primer assay for another known azole-resistance-inducing mutation, *ERG11* (G1885A). We used g-Block DNA templates for each mutation DNA region and combined varying concentrations (10^6^, 10^4^, and 10^3^ copies) of *ERG11*(G1885A) with 10^6^ copies of *ERG11*(A434T) in a duplex reaction. For the duplex reaction, we added *ERG11*(A434T) SuperSelective primer CX2475 and the reverse primer CX2477 along with *ERG11*(G1885A) SuperSelective primer CX2607 (20-14/15-6:1:0) and the reverse primer CX2328. As expected, each primer set specifically detected its intended mutant target without loss of specificity. *ERG11*(G1885A) generated a fluorescent signal in the Cal Fluor Red 610 channel with 10^6^, 10^4^, and 10^3^ copies of the DNA template, while *ERG11*(A434T) was simultaneously detected in the FAM channel with 10^6^ copies of the DNA template in the same reaction (Fig 4A and 4B). Importantly, the assay detected as few as 10^3^ copies of *ERG11*(G1885A) in the presence of 10^6^ copies of *ERG11*(A434T), demonstrating the sensitivity of the duplex format (Fig 4B). These results demonstrate that the SSP-PCR platform can selectively and sensitively detect two resistance-associated SNPs in a single tube reaction, highlighting its potential as a multiplex assay for high-throughput screening.

**Figure 4.**
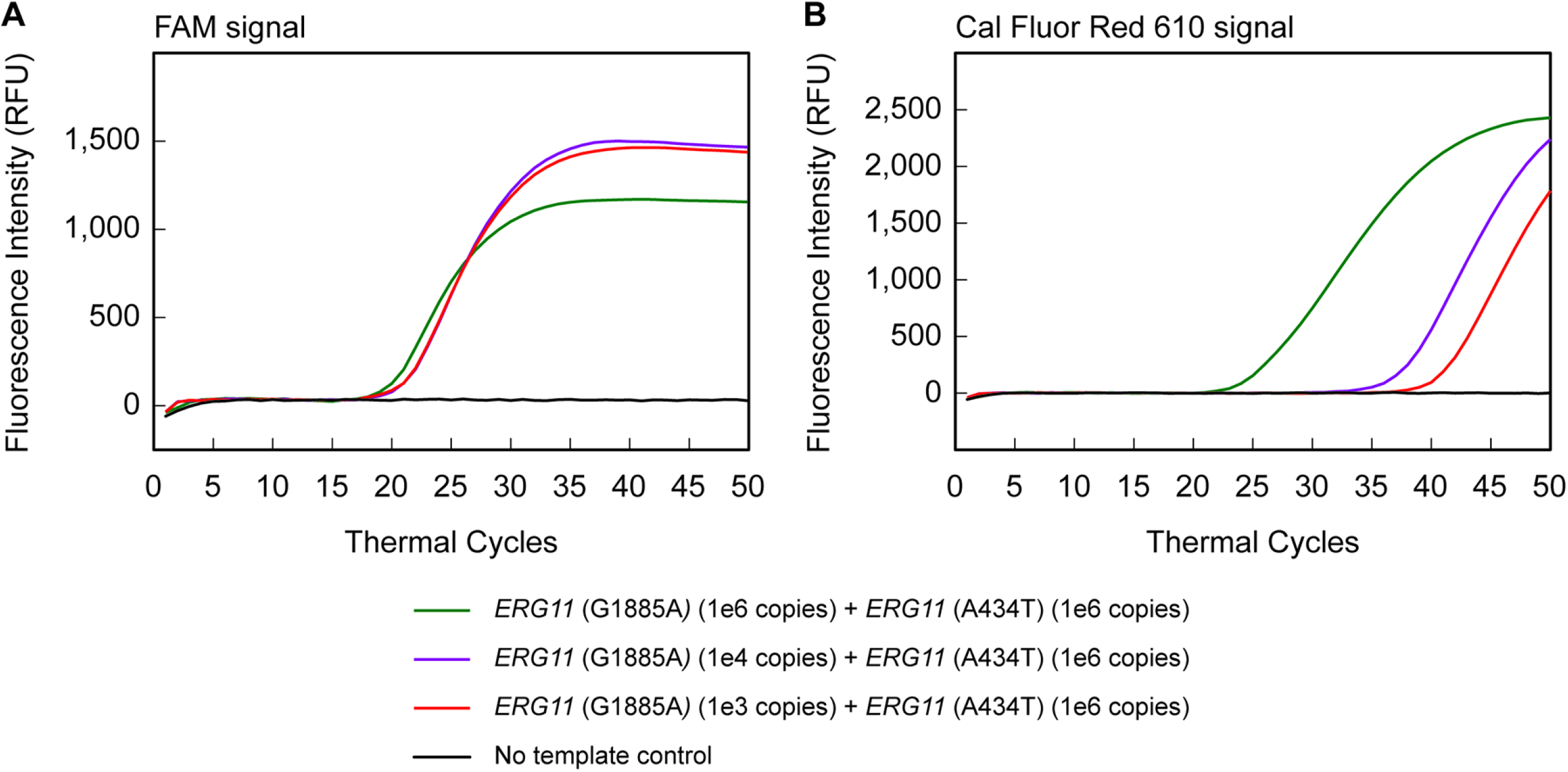
Concurrent detection of *ERG11*(A434T) and *ERG11*(G1885A) mutations in a duplex reaction using SSP-PCR. A duplex SSP-PCR assay simultaneously detects two azole resistance-associated SNPs – *ERG11*(A434T) and *ERG11*(G1885A) in two different fluorophore channels. Varying concentrations (10^6^, 10^4,^ and 10^3^ copies) of *ERG11*(G1885A) templates were combined with 10^6^ copies of the *ERG11*(A434T) template. (A) FAM-channel amplification curves showing specific detection of the *ERG11*(A434T) target. (B) Cal Fluor Red 610-channel amplification curves showing specific detection of *ERG11*(G1885A) target. Each SuperSelective primer-probe set selectively detected its intended target.

## Discussion

Cryptococcosis is one of the leading causes of death in immunocompromised patients, imposing a disproportionate public health burden in resource-limited settings. Early diagnosis is critical to guide proper therapeutic interventions to reduce cryptococcosis-associated morbidity and mortality (5). However, the current diagnosis for cryptococcosis is inadequate. Even in a closely monitored high-risk population, confirmation of cryptococcal meningitis takes 2-3 weeks from the onset of symptoms (23). Diagnosis is mainly based on a combination of histopathology, direct microscopy, fungal culture, and/or a positive *Cryptococcus* antigen test (24). Antigen-based assays, like detection of polysaccharide capsule in serum and cerebrospinal fluid by enzyme-linked immunosorbent assays (ELISA), latex flow assay (LFA), or Latex agglutination, are highly sensitive yet less effective for detection of pulmonary infection (5), is the current clinical gold standard for cryptococcal diagnosis. Moreover, PCR-based methods have been developed to either identify all *Cryptococcus* species based on fungal internal transcribed spacer (*ITS*) region or distinguish different species with species specific primers targeting the fungal mitochondrial cytochrome b gene (cyt b) sequence (25)(26). However, none of these currently available diagnostic methods can identify cryptococcal antifungal drug resistance, including heteroresistance in clinical setting.

Heteroresistance in *Cn* is mainly caused by chromosomal duplication resulting in aneuploidy and is most frequently observed in patients with cryptococcal meningitis undergoing FLC monotherapy (8)(27)(28)(29). Rapid identification of antifungal resistance remains a significant diagnostic challenge in pulmonary cryptococcosis, particularly when the resistance arises from SNPs within genetically heterogeneous or low-burden clinical specimens. Although sequencing-based approaches, such as next-generation sequencing, can detect SNPs, they are expensive, time-consuming, and require specialized equipment and experienced personnel. Conventional PCR assays with differently colored TaqMan probes or molecular beacons can be used to distinguish mutants from wild type. However, these assays utilize outside primers that amplify both mutant and their closely related wild type sequences, resulting in weak amplification of the mutant target and limiting the ability to detect rare mutant sequences in the background of abundant wild type DNA(17)(19). In hopes of overcoming these limitations, SSP-PCR assays capable of discriminating the single base changes in mixed genetic backgrounds without amplifying the wild type template offer a promising alternative. (17)(19).

In this study, we developed an SSP-PCR platform capable of detecting two known azole resistance-associated single-nucleotide polymorphism in *ERG11* gene of *C. neoformans*. The ability to detect the *ERG11*(A434T) mutant allele in synthetic g-Block templates despite the presence of wild-type sequence underscores the single-base specificity of this assay. This performance extended to genomic DNA from a hyper-resistant isolate *mrl1* that contains this mutation and, importantly, to samples derived from a murine infection model in which mice were infected with either H99, *mrl1*, or a combination of H99 and *mrl1*. In the mixed-infection setting, the platform successfully identified the *mrl1*-associated mutation, demonstrating its ability to detect mutant alleles in the presence of an abundant wild-type background in vivo. Additionally, coupling SuperSelective primers with molecular beacon probes enabled simultaneous detection of two azole resistance-associated mutations, *ERG11*(A434T) and *ERG11*(G1885A), in a single reaction. These findings highlight the promise of this platform as a rapid molecular diagnostic strategy for detecting resistance-associated alleles, particularly in heterogeneous samples containing both susceptible and resistant populations.

SSP-PCR assays, by virtue of their design (Fig.1A), exponentially amplify the mutant DNA template even in the presence of their closely related wild-type DNA template. During the PCR annealing, the SuperSelective primer’s foot sequence forms a stable foot-target hybrids that persist for longer times, which is sufficient to permit a polymerase to initiate the synthesis of an amplicon, while the mismatched foot-target hybrids are so short-lived that they very rarely initiate the synthesis of an amplicon. Since the foot sequence is very short, it results in a large enthalpic difference in the binding of the primer to the perfectly complementary mutant target sequence compared to the binding to the mismatched wild-type target. Adding tetramethylammonium chloride (TMAC) in the PCR assay further enhances this by stabilizing the matched mutant hybrids while destabilizing the mismatched wild-type target (30). Combining the specificity of SuperSelective primers with molecular beacons results in a versatile diagnostic platform that has been used to detect drug resistance-associated mutations in several different infectious diseases and cancer (17, 18). A clear precedent of the SSP-PCR assay demonstrated whether a patient is infected with *Mycobacterium tuberculosis*, the bacterial load, and whether the strain is resistant to the antibiotic rifampin, all within two hours using the patient’s sputum (31). A recent study demonstrated an alternative design for SuperSelective primers known as Self-Competitive Fishing primer that combined SuperSelective primers with wild-type blocking oligonucleotides to prevent amplification of wild-type DNA while enhancing mutant DNA detection in endometrial cancer, offering a practical alternative to next-generation sequencing (32).

MBs are added prior to amplification, they fluoresce in real-time in the presence of their targets, enabling quantitative measurement of the starting template while minimizing background noise due to primer dimers that commonly affect PCR signal specificity. MBs have been employed to detect *ERG11* mutations in *Candida auris* (33), and FKS-associated echinocandin resistance in *Candida auris* (33), *C. albicans* (34), and *C. glabrata* (35). However, a limitation of MBs or other hybridization probe-based SNP detection methods is that they cannot detect a rare mutant population when it’s less than 5-10% of the total population. When combined with SuperSelective primers, this configuration enhances SNP discrimination within mixed-allele pools. Thus, their structural and thermodynamic properties make them suitable candidates for SNP-level diagnostics (17)(30)(31)(18)(19).

Despite these strengths, our current study has a few limitations. For example, unlike bedside cryptococcal antigen detection LFA assays, our platform requires genomic DNA isolation, which might limit immediate point-of-care implementation. But this limitation could be alleviated by developing quick DNA isolation assays. Our PCR validation was performed under experimentally heterogeneous conditions rather than large-scale clinical cohorts, and further validation using clinical samples is necessary to establish diagnostic sensitivity and specificity. Additionally, the current assay targets only a few resistance-associated mutations, whereas clinical resistance may arise from multiple mutations within *ERG11* or other loci. However, the combination of SuperSelective primers with MB technology enables us to design multiple mutant-specific SuperSelective primers with different fluorophores for a multiplex panel in a single tube. Therefore, our future work will focus on developing a multiplex panel capable of simultaneously detecting multiple azole resistance mutations within a single reaction, thereby enhancing the clinical utility and scalability.

In conclusion, we have established a platform that combines mutation-specific amplification with fluorescent probing for rapid SNP-based detection of azole resistance in *Cryptococci*. To our knowledge, this is the first molecular assay to distinguish mutant targets under experimentally heterogeneous conditions without cross-reactivity, even *in in* vivo-infected tissues. With further development into a multiplex panel, this platform may complement existing diagnostic strategies to improve clinical outcomes.

## Materials and Methods

### Strains and growth conditions

All fungal strains were streaked onto YPD agar plates and grown in YPD liquid medium at 30 °C overnight with shaking. The following fungal strains were used in the study: *C. neoformans* reference strain H99 and *C. neoformans* isolate carrying *ERG11* Y145F (A434T) mutation (*mrl)*, a clinical isolate gifted by Dr. Jun Kwon-Chung from NIH. Chemicals and primers were purchased from Sigma-Aldrich (St Louis, MO) unless specified.

### Isolation of genomic DNA from fungi

Fungal cells were scraped from YPD plates and transferred to a 2 mL screw cap tube with 0.6 mL of TENTS buffer (10 mM Tris-HCl, pH 7.5; 1 mM EDTA, pH 8; 100 mM NaCl; 2% Trinton X-100; 1% SDS) and 0.4 mL of acid-wash beads. Bead-beat for 6 mins on a tabletop bead shaker. 0.5 mL of phenol: chloroform: isoamyl alcohol was added, and the tubes were left open for 5 mins. Cells were then centrifuged for 15 mins at 10,000 rpm. The aqueous phase was transferred into a fresh tube, and 1/10^th^ volume of 3 M Sodium acetate and 2.5 volumes of 100% EtOH were added. DNA was precipitated at -20°C for 30 mins. Cells were centrifuged for 15 mins at 10,000 rpm, and the pellet was washed with 1 mL of 70% EtOH and centrifuged for 5 mins. Supernatants were discarded, and the tubes were kept upside down on a clean paper towel to allow the pellets to air dry. The pellets were dissolved in 100 *µ*L of autoclaved ddH_2_O.

### Mouse infection

Each group of three C57BL/6 mice (Jackson Laboratory, Bar Harbor, ME) were intranasally infected with 10^5^ yeast cells per mouse of *C. neoformans* H99, the *mrl* strain containing *ERG11*(A434T) mutation or in a 1:1 combination (*ERG11*(A434T): *ERG11*WT), respectively. Mice were sacrificed 7 days post-infection, and lungs were harvested for the SSP-PCR assay. The mouse lungs were homogenized in 1xPBS using a homogenizer, and total DNA was extracted using the CTAB method as described below.

### Isolation of genomic DNA from infected mouse lung using CTAB method

Briefly, 0.6 mL of lung homogenate was transferred into the 2 mL screwcap tubes, and 0.7 mL of autoclaved ddH_2_O was added. The tubes were placed at 4°C overnight to lyse the mammalian cells. The next day, the tubes were centrifuged at 7,000 rpm for 10 mins, and the supernatant was discarded. The cells were washed again with autoclaved ddH_2_O. The samples were resuspended in Urea buffer (0.5 M NaCl, 20 mM Tris, 20 mM EDTA, and 2% SDS) for 4 h at room temperature with gentle agitation. Cell pellets collected by 5 mins centrifugation at 2,000 rpm and 0.4 mL of acid-wash beads were added to the tube along with the 0.4 mL of CTAB buffer (1 M Tris pH 7.5, 5 M NaCl, 0.5 M EDTA, CTAB (Sigma-Aldrich (St Louis, MO) M7635), beta-mercaptoethanol (14 M)), followed by bead-beat for 3-5 mins. Additional CTAB buffer was added to reach ∼ 600 µL, and the tubes were incubated at 65°C on a heat block before cooling to room temperature. An equal amount of chloroform was added, gently mixed, and centrifuged at 8,000 rpm for 10 mins. The clear aqueous phase was transferred to a fresh tube; an equal volume of isopropanol was added. Incubated 30 mins at -20°C, followed by centrifugation at 10,000 rpm for 10 mins. Supernatant was discarded. The pellet was washed with 70% ethanol and air-dried. The pellet was dissolved in TE buffer.

### Primers, g-Blocks and molecular beacons

SuperSelective primer sequences were examined with the Mfold web server and the OligoAnalyzer computer program (Integrated DNA Technologies, Coralville, IA) to ensure that they do not form hairpin structures, self-dimers, or heterodimers with conventional reverse primers. The primers were purchased from Integrated DNA Technologies or Sigma-Aldrich (St Louis, MO), and the MB probes for detecting the amplicons were purchased from Biosearch Technologies (Petaluma, CA). The synthetic g-Block DNAs of *C. neoformans* azole-associated *ERG11* mutations: Y145F (A434T) and G484S (G1885A) were purchased from Integrated DNA Technologies (Table S1).

### Real-time PCR assays

Monoplex real-time PCR assays were performed in 25 *µ*L volumes containing 200 nM of each forward (SuperSelective or conventional) and reverse primer, 1.5 mM MgCl_2_, 20 mM of TMAC, Taq 2x MasterMix (Platinum Hot start PCR 2X) and 1x SYBR Green were purchased from ThermoFischer Scientific (Waltham, MA) and added to monitor template amplification (20 *µ*L reaction mix + 5 *µ*L of template).

Duplex real-time PCR assays were carried out in 25 *µ*L volumes containing 200 nM of each forward (SuperSelective or conventional) and reverse primer, 1.5 mM MgCl2 (Invitrogen), 20 mM of TMAC, Taq 2x MM (Platinum Hot start PCR 2X) and 200 nM of each MB for monitoring amplicon generation during the annealing stage of each thermal cycle (20 *µ*L reaction mix + 5 *µ*L of template).

All amplifications were carried out in 200 *µ*L white polypropylene PCR tubes (USA Scientific, Ocala, FL) in a Bio-Rad CFX96 detection system (Bio-Rad Laboratories, Hercules, CA). All reactions were carried out using the following PCR cycle details: Hot start 95°C for 2 mins to activate the polymerase, followed by 50 repeats of a 95°C 15 seconds denaturation step and 60°C for 20 seconds annealing step, while monitoring the fluorescence.

## Ethics Statement

Animal studies were performed at Rutgers University Newark campus animal facility. All studies were conducted following biosafety level 2 (BSL-2) protocols and procedures approved by the Institutional Animal Care and Use Committee (IACUC) and Institutional Biosafety Committee of Rutgers University under protocol 999901066. Animal studies were compliant with all applicable provisions established by the Animal Welfare Act and the Public Health Services (PHS) Policy on the Humane Care and Use of Laboratory Animals.

## Data Availability

Data are provided within the manuscript or supplemental material. The strains and other materials for the current study are available from the corresponding authors on reasonable request.

## Author contributions

SP, data curation, formal analysis, investigation, methodology, and writing-original draft and editing | HHX and SW, data curation & methodology | SM and CX, conceptualization, formal analysis, supervision, funding acquisition, project administration, resources & writing – review and editing.

## Acknowledgements

We thank all members of Xue and Marras Laboratories for their support. We thank Dr. Jun Kwon-Chung for providing the *mrl1* clinical isolate. The Xue laboratory is supported by NIH grants AI155647, AI123315, AI169769, and AI141368. This work is submitted in partial fulfillment of the PhD degree in Biomedical Sciences at Rutgers, The State University of New Jersey, USA.

## References

1. Denning DW. 2024. Global incidence and mortality of severe fungal disease. Lancet Infect Dis 24:e428–e438.

2. Kainz K, Bauer MA, Madeo F, Carmona-Gutierrez D. 2020. Fungal infections in humans: the silent crisis. Microb Cell 7:143–145.

3. Rayens E, Norris KA. 2022. Prevalence and Healthcare Burden of Fungal Infections in the United States, 2018. Open Forum Infect Dis 9:ofab593.

4. Park BJ, Wannemuehler KA, Marston BJ, Govender N, Pappas PG, Chiller TM. 2009. Estimation of the current global burden of cryptococcal meningitis among persons living with HIV/AIDS. Aids 23:525–30.

5. McHale TC, Boulware DR, Kasibante J, Ssebambulidde K, Skipper CP, Abassi M. 2023. Diagnosis and management of cryptococcal meningitis in HIV-infected adults. Clin Microbiol Rev 36:e0015622.

6. Hope W, Stone NRH, Johnson A, McEntee L, Farrington N, Santoro-Castelazo A, Liu X, Lucaci A, Hughes M, Oliver JD, Giamberardino C, Mfinanga S, Harrison TS, Perfect JR, Bicanic T. 2019. Fluconazole Monotherapy Is a Suboptimal Option for Initial Treatment of Cryptococcal Meningitis Because of Emergence of Resistance. mBio 10:e02575–19

7. Roemer T, Krysan DJ. 2014. Antifungal drug development: challenges, unmet clinical needs, and new approaches. Cold Spring Harb Perspect Med 4:a019703

8. Mondon P, Petter R, Amalfitano G, Luzzati R, Concia E, Polacheck I, Kwon-Chung KJ. 1999. Heteroresistance to fluconazole and voriconazole in Cryptococcus neoformans. Antimicrob Agents Chemother 43:1856–61.

9. Sionov E, Chang YC, Garraffo HM, Dolan MA, Ghannoum MA, Kwon-Chung KJ. 2012. Identification of a Cryptococcus neoformans cytochrome P450 lanosterol 14α-demethylase (Erg11) residue critical for differential susceptibility between fluconazole/voriconazole and itraconazole/posaconazole. Antimicrob Agents Chemother 56:1162–9.

10. Sionov E, Lee H, Chang YC, Kwon-Chung KJ. 2010. Cryptococcus neoformans overcomes stress of azole drugs by formation of disomy in specific multiple chromosomes. PLoS Pathog 6:e1000848.

11. Varma A, Kwon-Chung KJ. 2010. Heteroresistance of Cryptococcus gattii to fluconazole. Antimicrob Agents Chemother 54:2303–11.

12. Sionov E, Chang YC, Kwon-Chung KJ. 2013. Azole heteroresistance in Cryptococcus neoformans: emergence of resistant clones with chromosomal disomy in the mouse brain during fluconazole treatment. Antimicrob Agents Chemother 57:5127–30.

13. Binnicker MJ, Jespersen DJ, Bestrom JE, Rollins LO. 2012. Comparison of Four Assays for the Detection of Cryptococcal Antigen. Clinical and Vaccine Immunology 19:1988– 1990.

14. Bosco-Borgeat ME, Mazza M, Taverna CG, Córdoba S, Murisengo OA, Vivot W, Davel G. 2016. Amino acid substitution in Cryptococcus neoformans lanosterol 14-α-demethylase involved in fluconazole resistance in clinical isolates. Rev Argent Microbiol 48:137–42.

15. Rodero L, Mellado E, Rodriguez AC, Salve A, Guelfand L, Cahn P, Cuenca-Estrella M, Davel G, Rodriguez-Tudela JL. 2003. G484S amino acid substitution in lanosterol 14-alpha demethylase (ERG11) is related to fluconazole resistance in a recurrent Cryptococcus neoformans clinical isolate. Antimicrob Agents Chemother 47:3653–6.

16. Kano R, Okubo M, Hasegawa A, Kamata H. 2017. Multi-azole-resistant strains of Cryptococcus neoformans var. grubii isolated from a FLZ-resistant strain by culturing in medium containing voriconazole. Med Mycol 55:877–882.

17. Vargas DY, Kramer FR, Tyagi S, Marras SAE. 2016. Multiplex Real-Time PCR Assays that Measure the Abundance of Extremely Rare Mutations Associated with Cancer. PLOS ONE 11:e0156546.

18. Kramer FR, Vargas DY. 2021. SuperSelective primer pairs for sensitive detection of rare somatic mutations. Scientific Reports 11:22384.

19. Vargas DY, Tyagi S, Marras SAE, Moerzinger P, Abin-Carriquiry JA, Cuello M, Rodriguez C, Martinez A, Makhnin A, Farina A, Patel C, Chuang TL, Li BT, Kramer FR. 2022. Multiplex SuperSelective PCR Assays for the Detection and Quantitation of Rare Somatic Mutations in Liquid Biopsies. J Mol Diagn 24:189–204.

20. Tyagi S, Kramer FR. 1996. Molecular beacons: probes that fluoresce upon hybridization. Nature biotechnology 14:303–308.

21. Tyagi S, Bratu DP, Kramer FR. 1998. Multicolor molecular beacons for allele discrimination. Nature biotechnology 16:49–53.

22. Tyagi S, Kramer FR. 2012. Molecular beacons in diagnostics. F1000 Med Rep 4:10.

23. Perfect JR, Bicanic T. 2015. Cryptococcosis diagnosis and treatment: What do we know now. Fungal Genet Biol 78:49–54.

24. Chang CC, Hall V, Cooper C, Grigoriadis G, Beardsley J, Sorrell TC, Heath CH. 2021. Consensus guidelines for the diagnosis and management of cryptococcosis and rare yeast infections in the haematology/oncology setting, 2021. Intern Med J 51 Suppl 7:118–142.

25. Lau A, Sorrell TC, Chen S, Stanley K, Iredell J, Halliday C. 2008. Multiplex tandem PCR: a novel platform for rapid detection and identification of fungal pathogens from blood culture specimens. J Clin Microbiol 46:3021–7.

26. Tay E, Chen SC, Green W, Lopez R, Halliday CL. 2022. Development of a Real-Time PCR Assay to Identify and Distinguish between Cryptococcus neoformans and Cryptococcus gattii Species Complexes. J Fungi (Basel) 8:462.

27. Stone NR, Rhodes J, Fisher MC, Mfinanga S, Kivuyo S, Rugemalila J, Segal ES, Needleman L, Molloy SF, Kwon-Chung J, Harrison TS, Hope W, Berman J, Bicanic T. 2019. Dynamic ploidy changes drive fluconazole resistance in human cryptococcal meningitis. J Clin Invest 129:999–1014.

28. Billmyre RB, Clancey SA, Heitman J. 2017. Natural mismatch repair mutations mediate phenotypic diversity and drug resistance in Cryptococcus deuterogattii. Elife 6:e28802.

29. Boyce KJ, Wang Y, Verma S, Shakya VPS, Xue C, Idnurm A. 2017. Mismatch Repair of DNA Replication Errors Contributes to Microevolution in the Pathogenic Fungus Cryptococcus neoformans. mBio 8:e00595–17.

30. Vargas DY, Marras SAE, Tyagi S, Kramer FR. 2018. Suppression of Wild-Type Amplification by Selectivity Enhancing Agents in PCR Assays that Utilize SuperSelective Primers for the Detection of Rare Somatic Mutations. J Mol Diagn 20:415–427.

31. Narang A, Marras SAE, Kurepina N, Chauhan V, Shashkina E, Kreiswirth B, Varma-Basil M, Vinnard C, Subbian S. 2022. Ultrasensitive Detection of Multidrug-Resistant Mycobacterium tuberculosis Using SuperSelective Primer-Based Real-Time PCR Assays. Int J Mol Sci 23:15752.

32. Wu C-C, Hsiao Y-C, Lin Z-Y, Chiu P-H, Chang C-L. 2026. A Novel Self-Competitive Fishing Primer qPCR Approach for Efficient POLE Mutation Detection in Endometrial Cancer Molecular Classification. Current Issues in Molecular Biology 48:257.

33. Hou X, Lee A, Jiménez-Ortigosa C, Kordalewska M, Perlin DS, Zhao Y. 2019. Rapid Detection of ERG11-Associated Azole Resistance and FKS-Associated Echinocandin Resistance in Candida auris. Antimicrob Agents Chemother 63:e01811–18.

34. Balashov SV, Park S, Perlin DS. 2006. Assessing resistance to the echinocandin antifungal drug caspofungin in Candida albicans by profiling mutations in FKS1. Antimicrob Agents Chemother 50:2058–63.

35. Zhao Y, Nagasaki Y, Kordalewska M, Press EG, Shields RK, Nguyen MH, Clancy CJ, Perlin DS. 2016. Rapid Detection of FKS-Associated Echinocandin Resistance in Candida glabrata. Antimicrob Agents Chemother 60:6573–6577.

